# Chloroquine Kills Hair Cells in Zebrafish Lateral Line and Murine Cochlear Cultures: Implications for Ototoxicity

**DOI:** 10.1101/2020.04.14.041731

**Authors:** Samantha N. Davis, Patricia Wu, Esra D. Camci, Julian A. Simon, Edwin W Rubel, David W. Raible

**Affiliations:** Virginial Merrill Bloedel Hearing Research Center, University of Washington Seattle, WA, USA; Department of Speech and Hearing Sciences, University of Washington Seattle, WA, USA; Department of Biological Structure, University of Washington Seattle, WA, USA; Department of Otolaryngology – Head and Neck Surgery, University of Washington Seattle, WA, USA; Fred Hutch Cancer Research Center Seattle, WA, USA

**Keywords:** ototoxicity, anti-malarial, hearing loss, vestibular loss

## Abstract

Hearing and balance deficits have been reported during and following treatment with the antimalarial drug chloroquine. However, experimental work examining the direct actions of chloroquine on mechanoreceptive hair cells in common experimental models is lacking. This study examines the effects of chloroquine on hair cells using two common experimental models: the zebrafish lateral line and neonatal mouse cochlear cultures. Zebrafish larvae were exposed to varying concentrations of chloroquine phosphate or hydroxychloroquine for 1 hr or 24 hr, and hair cells assessed by antibody staining. A significant, dose-dependent reduction in the number of surviving hair cells was seen across conditions for both exposure periods. Hydroxycholroquine showed similar toxicity. In mouse cochlear cultures, chloroquine damage was specific to outer hair cells in tissue from the cochlear basal turn, consistent with susceptibility to other ototoxic agents. These findings suggest a need for future studies employing hearing and balance monitoring during exposure to chloroquine and related compounds, particularly with interest in these compounds as therapeutics against viral infections including coronavirus.

## 1. Introduction

Quinoline antimalarial drugs have long been implicated as a cause of hearing loss, tinnitus, dizziness, and severe imbalance (Bernard, 1985; Hart and Naunton, 1964; Matz and Naunton, 1968; Mukherjee, 1979). This side effect profile prompted the transition from quinine to chloroquine, and further development of hydroxychloroquine (Matz and Naunton, 1968; Scherbel et al., 1958; Shrivastav et al., 2016). Chloroquine interferes with parasite growth in red blood cells, thereby protecting against or disrupting the progression of malaria (Hoppe et al., 2004; Kapishnikov et al., 2019). Despite the development of new drugs for the prevention and treatment of malaria, their ototoxic side effects have persisted.

Chloroquine and hydroxychloroquine have many uses in addition to being anti-malarial agents, and may therefore cause adverse events in multiple patient populations. Chloroquine or hydroxychloroquine are also commonly prescribed to treat autoimmune and collagen conditions such as rheumatoid arthritis, scleroderma, and systemic lupus erythematosus, as well as amebiasis (Bernard, 1985; Matz and Naunton, 1968) and other off-label applications (Al-Bari, 2015; Bogaczewicz and Sobów, 2017; Plantone and Koudriavtseva, 2018); these patients are more likely to experience ear-related symptoms due to the higher dosing and prolonged exposure to chloroquine. Most recently chloroquine has been promoted as a therapeutic against coronavirus disease COVID-19 (Cortegiani et al., 2020; Sahraei et al., 2020; Zhou et al., 2020).

Patients receiving chloroquine may experience sensorineural hearing loss—as extreme as a complete loss of mid- and high-frequency hearing—(Mukherjee, 1979), tinnitus (Bernard, 1985), and vestibular deficits such as vestibular paresis (Matz and Naunton, 1968; Mukherjee, 1979), as quickly as two hours after an injection of chloroquine (Mukherjee and Mukherjee, 1979). These side effects are all associated with the inner ear sensory structures, raising the suspicion of objective ototoxicity. Hydroxychloroquine has been associated with similar side effects, with over 700 reported adverse events linked to the ear and labyrinthine structure in the United States in the last 5 years (U.S. Food and Drug Administration, 2019).

Unfortunately, knowledge of adverse effects has largely come from case studies, leaving the mechanisms and prevalence of these effects to be unclear. Both reversible (Bernard, 1985; Mukherjee, 1979) and irreversible (Bernard, 1985; Matz and Naunton, 1968; Toone et al., 1965) symptoms concerning hearing and balance have been reported, but there is little understanding how these antimalarial drugs affect the structures of the inner ear and central nervous system involved in hearing and balance. Additionally, there is a lack of animal studies exploring the direct effects of chloroquine and hydroxychloroquine on the peripheral auditory system, and specifically, on mechanosensory hair cells. Here, we use the zebrafish lateral line and mouse neonatal cochlear cultures to test the effect of chloroquine and hydroxychloroquine on mechanosensory hair cell survival.

The zebrafish lateral line is a good model for understanding hair cell function and dysfunction (Harris et al., 2003; Ou et al., 2012; Pickett et al., 2018). The lateral line is a system of mechanosensory hair cells on the surface of body that allows fish to detect fluid displacement. Inputs from this system provide a way for fish to orient themselves in water for behaviors such as schooling and predator avoidance (Bak-Coleman et al., 2013; Thomas et al., 2015). The system consists of neuromasts, clusters of hair cells surrounded by support cells. These hair cells depolarize as their stereocilia are displaced and send signals to the brain with associated nerve fibers, similar to the functions of hair cells in the auditory and vestibular labyrinth. Lateral line hair cells are easily accessible for experimental manipulation and provide a rapid system for screening compounds for ototoxicity (Chiu et al., 2008; Hirose et al., 2011; Owens et al., 2008) as well as investigating the specific mechanisms of toxin entry into the cell and subsequent cell death (Hailey et al., 2017; Owens et al., 2008; Santos et al., 2006). In addition, the zebrafish lateral line system has been used to screen for small molecules that protect hair cells from damage, with translation to mammalian systems (Chowdhury et al., 2018; Kitcher et al., 2019). Previous work in this system has largely focused on aminoglycoside antibiotics and cisplatin, known ototoxins that cause irreversible hearing and balance disorders in human patients. We then compared the effect of chloroquine in lateral line hair cells, an *in vivo* system, to their effect on neonatal mouse cochlear cultures, an established *in vitro* model for hair cell toxicity (Kotecha and Richardson, 1994; Richardson and Russell, 1991) Other known ototoxins cause comparable loss of hair cells in both systems (Kirkwood et al., 2017; Kitcher et al., 2019).

Here, we report that chloroquine causes specific loss of mechanosensory hair cells in the zebrafish lateral line and in the cultured neonatal mouse cochlea. We find rapid, dose-dependent cell death due to exposure to these compounds. We suggest that chloroquine-associated hearing loss and vestibular impairment in human patients may be due to loss of hair cells and warrants further study in vivo in mature mammals.

## 2. Methods

Procedures have been approved by the University of Washington Animal Care and Use Committee.

### 2.1 Zebrafish studies

Adult zebrafish (*Danio rerio*) were paired to produce embryos in the University of Washington fish facility. Embryos raised at 50 embryos per 100 mm2 petri dish in E2 embryo medium (14.97 mM NaCl, 500 μM KCl, 42 μM Na2HPO4, 150 μM KH2PO4, 1 mM CaCl2 dihydrate, 1 mM MgSO4, 0.714 mM NaHCO3, pH 7.2) and raised in an incubator at 28.5°C. At four days post-fertilization (dpf), larvae were fed live rotifers.

At 5–7 dpf, larvae were placed in 48-well plates with 10–12 larvae per well, each well containing a different concentration of chloroquine in 300 µL of embryo medium. This study used concentrations ranging from 25 to 1600 µM with either 1- or 24-hour exposure periods. Effects of hydroxychloroquine were investigated up to 400 µM.

Following exposure to the drug, zebrafish larvae were euthanized and then fixed in 4% paraformaldehyde overnight. Larvae were rinsed with phosphate-buffered saline (PBS) and then placed in blocking solution (1 % Triton-X 100, 5 % normal goat serum (NGS) in PBS) at room temperature for 1–2 hours. Zebrafish were then placed in an anti-parvalbumin antibody solution (monoclonal, 1:400 in 1% Triton-X, 1 % NGS, in PBS) at 4 °C overnight and then rinsed with 1% Triton-X 100 in PBS (PBS-T) thrice. Larvae were then placed in a secondary solution with Alexa 488 goat anti-mouse H+L fluorescent antibody (1:500, in 1 % Triton-X 100, 1 % NGS, in PBS) for 4 hours. Following the secondary antibody solution, zebrafish were rinsed with PBS-T and PBS, before being mounted with Fluoromount-G (Southern Biotech, Birmingham, AL, USA) between two coverslips (VWR Cat #84393-251). Hair cells were counted in four neuromasts (SO1, SO2, O1, and OC1) per fish using a Zeiss Axioplan II microscope at a total magnification of 200X (40X objective; N.A.=0.75) with FITC filtered fluorescent illumination.

### 2.2 Cochlear cultures

Cochlear cultures were prepared from C57BL/6J mice according to previously described protocols (Kitcher et al., 2019; Russell and Richardson, 1987). Explants were prepared from pups sacrificed at postnatal day 2 and plated on collagen covered coverslips with 50 µL of cochlear culture medium (93% DMEM-F12, 7% FBS and 10 μg/ml ampicillin). The culture and coverslip were incubated overnight in a Maximow slide assembly at 37°C.

Experiments were carried out in 2 mL total volume of modified culture media (98.6% DMEM-F12, 1.4% FBS and 10 μg/ml ampicillin) with varying concentration of chloroquine. After 24 hours of incubation, cultures were washed once with PBS, fixed in 4% paraformaldehyde for 1 hour at room temperature. Hair cells were labeled with 1:1000 Phalloidin conjugated to Alexa Fluor 568 (Invitrogen, A12380) and 1:1000 anti-Myosin-VIIa rabbit polyclonal antibody (Santa Cruz, sc-74516), 1:1000 anti-Sox2 polyclonal antibody overnight followed by 1:500 Alexa Fluor 488 and 1:500 Alexa Fluor 694 goat anti-rabbit IgG (Invitrogen, A-11034).

Tissue was imaged at the midpoint of the apical region and at the midpoint of the basal region of the cochlear coil using an Axioplan ll (Zeiss) upright microscope at a final magnification of 200x. MyosinVIIa labeled hair cell bodies and Sox2 labeled support cell nuclei were counted in a 100 µm length along the sensory epithelium of each microscope field. A statistical power analysis based on previous experiments in this system, with an alpha of 0.05 and power = 0.80 indicated that a minimum of n= 4 samples per group would be required (Rosner, 2011).

### 2.3 Data Analysis

Total hair cells remaining were compared to counts of the same neuromasts of untreated fish or untreated cochlear cultures to calculate the percentage of hair cells remaining. Hair cell survival was then related to concentration of chloroquine used in incubation. Data was analyzed by t-tests, and by one-way or two-way ANOVAs with posthoc comparisons using GraphPad Prism.

## 3. Results

To test the potential toxic effects of chloroquine, zebrafish were treated with differing concentrations of drug. Neuromasts displayed clearly visible hair cell death after treatment for 24h (Figure 1A,B), with increasing hair cell loss after exposure to increasing chloroquine concentrations during 24h exposure (Figure 1C). Significant differences in hair cell numbers were found between control conditions and concentrations of 100 µM and higher (100, 200, 400, 800, 1600) (p<0.05). Hair cell survival reached an apparent asymptote around 65% with concentrations of 400 µM and higher (400, 800, 1600). In subsequent experiments, concentrations were limited to 0–400 µM dosages. Figure 1D compares the effects of treating zebrafish with chloroquine for 1h and 24h. Two-way ANOVA with dose and exposure time as main factors yielded a highly significant main effect of concentration (F=12, p<0.0001), but insignificant effects of exposure time (F=3.06, p=0.0828) and a significant interaction term (F=4.5, p=0.0009). These results demonstrate that chloroquine exposure results in a rapid dose-dependent loss of zebrafish lateral line hair cells.

**Figure 1:**
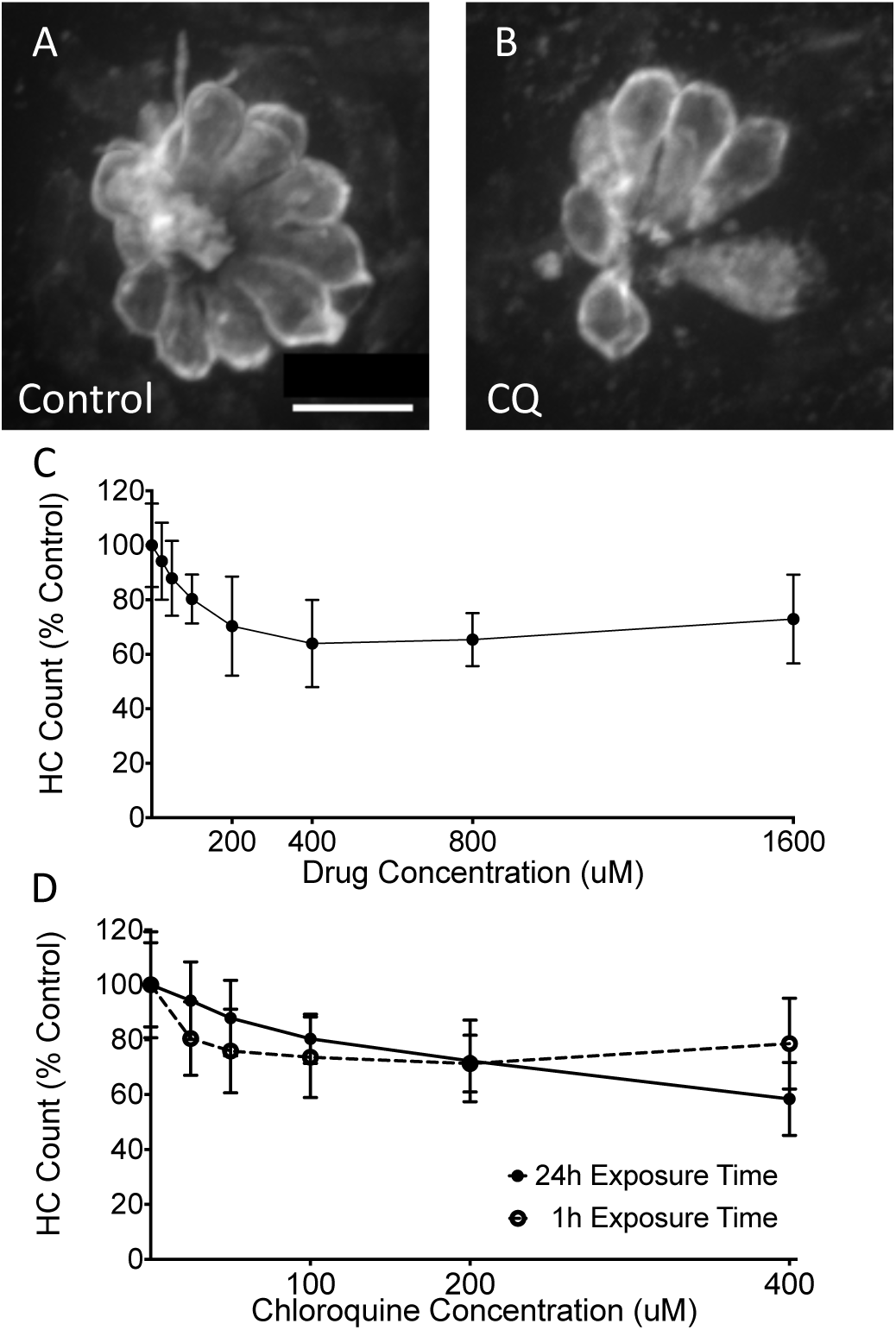
Chloroquine causes dose-dependent hair cell loss in the zebrafish lateral line system. Fluorescent imaging showed robust hair cell numbers in control neuromasts (A) and reduced numbers in fish exposed to 400 µM chloroquine for 24h (B). Hair cell counts show a decrease in hair cell viability as chloroquine concentration increased (F=12.28, p<0.0001;C). However, there was no significant change with concentrations from 400 µM to 1600 µM (p>0.05). (D) This pattern was consistent between 24- and 1-hour exposure times with no statistical significance (F=3.06, p = 0.0828). Error bars represent ±1 standard deviation. Scale bar = 10 µm.

As hydroxychloroquine is often used to replace chloroquine in clinical settings, these two drugs were compared for relative hair cell toxicity. Both drugs showed dose-dependent hair cell loss (Figure 2). Hydroxychloroquine showed slightly more toxicity than chloroquine (F=4.774, p=0.03; two-way ANOVA). These results demonstrate that both clinically-relevant chloroquine drugs show hair cell toxicity in the zebrafish model.

**Figure 2:**
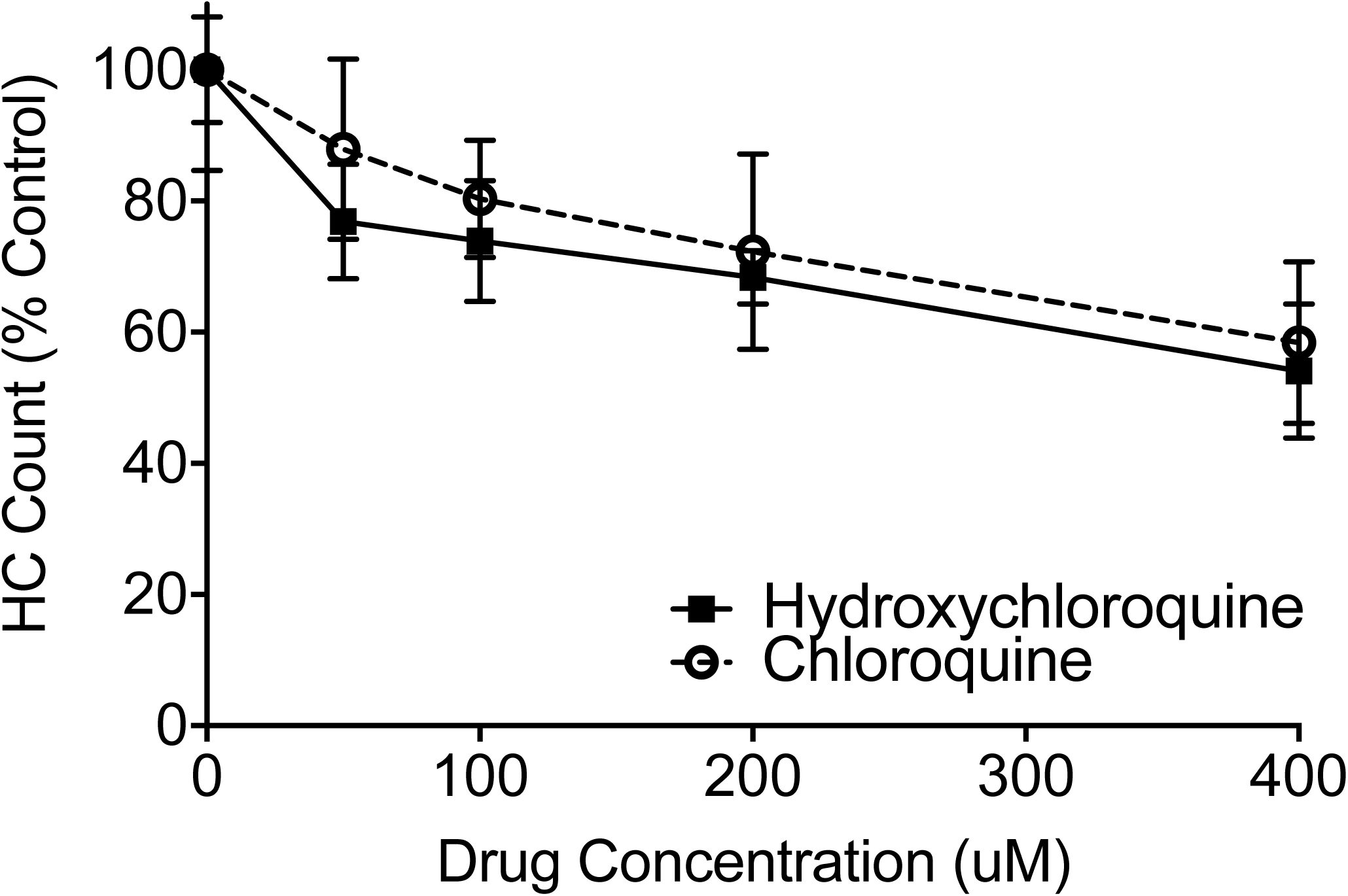
Hydroxychloroquine causes dose-dependent effects on hair cells in the zebrafish lateral line, similar to chloroquine. Dose-response functions show that hair cells exposed to hydroxychloroquine (solid line) survive at a similar rate as those exposed to chloroquine (dashed line) after a 24-hour treatment period. Two-way ANOVA demonstrate a small, but significant effect, with hydroxychloroquine being slightly more toxic (F=4.774, p=0.03). Error bars represent ±1 standard deviation.

Previous work has demonstrated that quinoline drugs, including chloroquine, can inhibit aminoglycoside ototoxicity by partially blocking mechanotransduction-dependent toxin uptake (Ou et al., 2012). We reasoned that the asymptotic nature of the chloroquine dose-response function might be due to its ability to block its own uptake. We performed two sets of experiments to test this idea. Function of the mechanotransduction channel was assessed through quantification of FM1-43X uptake after pre-treatment with chloroquine at doses that cause maximal hair cell death. Fish were also treated with benzamil, a MET channel blocker known to block drug uptake (Hailey et al., 2017; Rüsch et al., 1994) as a positive control (Figure 3). Chloroquine pre-treatment did not significantly alter FM1-43X uptake (p=0.2334; Dunn’s multiple comparison test), while benzamil-treated fish showed a highly significant decrease in FM1-43X uptake compared to controls (p<0.0001; Figure 3 A-D). We then examined whether chloroquine toxicity itself was altered by MET blockers (Figure 4). Neither 50 µM nor 200 µM benzamil co-treatment had a significant effect on chloroquine toxicity (p=0.9698 and 0.4943, respectively; Sidak’s multiple comparison test). Together these experiments suggest that the plateau of the dose-response function is not due to chloroquine effects on mechanotransduction.

**Figure 3:**
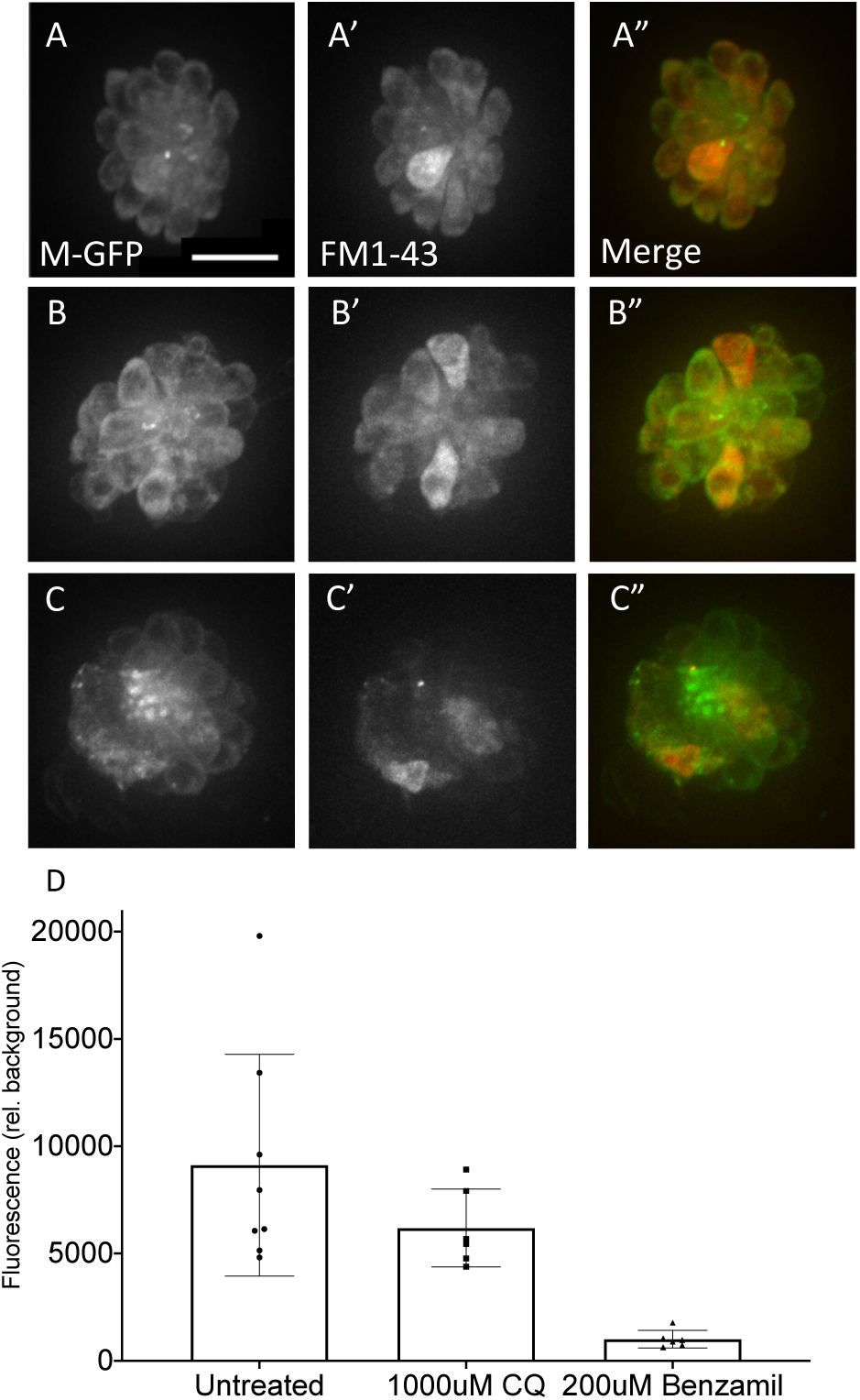
Chloroquine does not reduce MET channel activity. Fluorescent imaging showed no reduction of FM1-43-X uptake with pre-exposure to chloroquine (A–C). Brn3c fish, that have green fluorescent membrane markers, had no pre-treatment (A–A”), 1000 µM chloroquine for 15 minutes (B–B”), or 200 µM benzamil for 15 minutes (C–C”). Dunn’s multiple comparisons confirmed that red fluorescence is not significantly attenuated in fish pre-treated with a high dose of chloroquine (p=0.2334), while uptake is significantly lower in those fish treated with benzamil (p<0.0001), a known MET channel blocker (D; n=6–8 fish per condition). Error bars represent ±1 standard deviation. Scale bar = 10 µm

**Figure 4:**
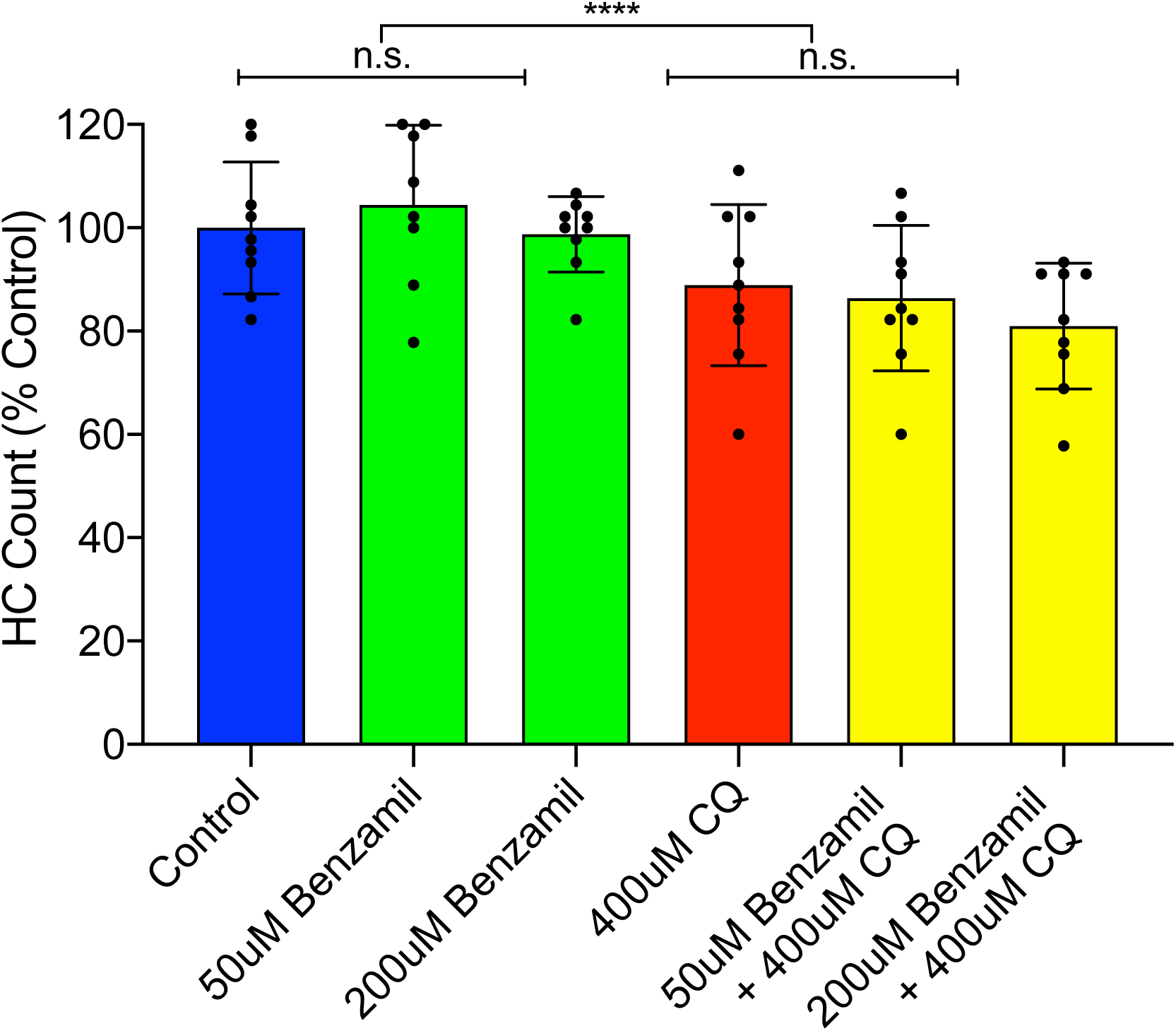
Chloroquine-induced cell death is not affected by the presence of a MET channel blocker. Co-treatment of chloroquine and benzamil showed no significant change in hair cell viability, though chloroquine exposure did have a small but significant effect on hair cell survival (p<0.0001). Error bars represent ±1 standard deviation.

To test whether mammalian hair cells degenerate in response to chloroquine, neonatal mouse cochlear cultures were exposed to 0–200 µM chloroquine for 24 hours (Figure 5). We compared responses of cultures from apical (Figure 5B) and basal (Figure 5C) regions of the cochlea. Apical regions showed no significant inner or outer hair cell loss with increasing concentrations of chloroquine (F=0.91, p=0.45; two-way ANOVA) Basal areas showed increasing hair cells loss with increasing chloroquine exposure (F=27.43, p <0.0001; two-way ANOVA). As for other ototoxins, basal outer hair cells were significantly more sensitive to chloroquine than inner hair cells (F=26.11, p <0.0001; two-way ANOVA). Analyses of the basal coil revealed that outer hair cells were significantly damaged (p<0.001) with 50% loss at 100 µM chloroquine exposure and almost complete loss was seen at 200 µM exposure, while there was a significant loss (p<0.05) of approximately 25% inner hair cells at the highest concentration. In order to evaluate any effects of chloroquine on supporting cells, Sox2-positive cells located between the inner hair cells and first row of outer hair cells were counted and compared to controls. A two-way ANOVA indicated no significant loss of chloroquine on supporting cell survival in either the apex or the base of cultures at any concentration (p>0.05). Together these results demonstrate that chloroquine damage at the concentrations tested is targeted to the hair cells and, like most other ototoxic drugs, most severely compromises basal turn outer hair cells as compared to inner hair cells.

**Figure 5:**
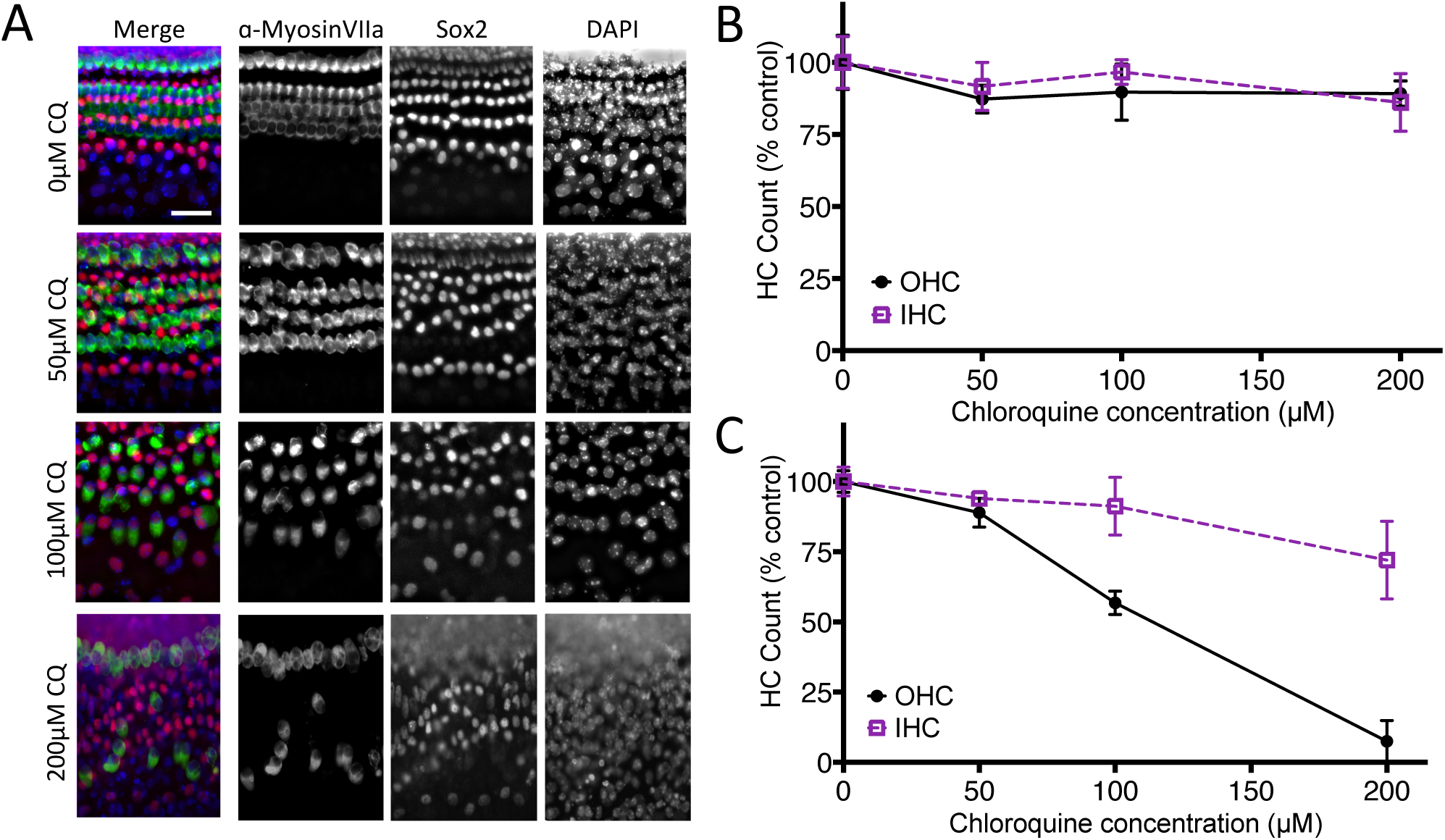
Chloroquine exposure causes dose-dependent hair cell loss in neonatal mouse organ of Corti cultures. (A) Fluorescent imaging of basal turn cultures treated with 0, 50, 100, and 200 µM chloroquine and labelled with antibodies specific to hair cells (Myosin Vlla), or organ of Corti supporting cells (Sox2), or all nuclei (DAPI). Note that 50 µM chloroquine exposure (row labeled 50 µM CQ) did not reveal clear loss or disruption of any cell type (compare with top row (0 µM CQ). However, 100 µM and 200 µM chloroquine exposure resulted in moderate and extreme loss of outer hair cells, respectively. Conversely, there was no obvious loss of other cell types at any treatment condition. (B) Apical coil cultures demonstrate little or no loss of inner or outer hair cells following treatment with chloroquine at any of the concentrations used (p=0.6087). (C) Basal coil hair cells, however, showed significantly decreased viability when treated with chloroquine (p<0.0001), and outer hair cells in the base were more dramatically affected by drug exposure than inner hair cells (F=26.11, p<0.0001). Error bars represent ±1 standard deviation. Scale bar = 20 µm

## 4. Discussion

This study describes dose-response analyses for the effects of chloroquine in the zebrafish anterior lateral line and in neonatal murine cochlear cultures. Chloroquine exposure was found to cause hair cell loss in both models. In zebrafish, variation in exposure time (1 hr vs 24 hr) did not result in significant differences in hair cell loss, implying that chloroquine uptake is rapid and affects hair cells soon after they are exposed. Exposure to hydroxychloroquine yielded comparable results to those from chloroquine exposure. For mouse cochlear cultures, reduction in hair cell counts were most dramatic in basal regions and in outer hair cells, consistent with differences in the sensitivity of hair cells to other ototoxins and reported hearing loss being limited to or most profound at high frequencies with most other ototoxins (Bielefeld et al., 2018; Chowdhury et al., 2018; Kotecha and Richardson, 1994). While there are no *in vivo* mammalian studies to compare with the results presented here, these results are consistent with the absence of hair cells with fetal exposure to chloroquine (Matz & Naunton, 1968).

Chloroquine and hydroxychloroquine have good oral bioavailability and reach high plasma levels in patients. Their pharmacokinetic profiles indicate extremely long elimination half-lives on (weeks to months) and large volumes of distribution indicating significant partitioning into tissues including cerebrospinal fluid (White 1985). Given these pharmacokinetic characteristics, it is entirely possible that drug concentration in the inner ear reach micromolar concentrations used in the experiments described here.

Because quinine users commonly reported similar side effects to that of patients taking chloroquine, it can be posited that the effects of these structurally-related quinoline drugs with the tissues of the inner ear are comparable. These results are consistent with reports of audiological and vestibular dysfunction after quinine treatment for malaria prophylaxis or treatment (Hennebert and Fernández, 1959; Karlsson et al., 1990; Nielsen-Abbring et al., 1990; Phillips-Howard and ter Kuile, 1995; Roche et al., 1990). Studies of quinine ototoxicity have shown extensive damage and deterioration of the stria vascularis and organ of Corti, especially to outer hair cells (Hennebert and Fernández, 1959). Exposed animals displayed minimal damage in the vestibular system, and impacted vestibular function recovered with time. Peripheral auditory function was reduced with quinine injections—acutely exposed animals improved fully or partially, and those receiving chronic dosing remained affected.

The mechanism by which chloroquine acts to kill hair cells is not yet known. The pharmacological mechanisms of action of chloroquine are subject of some controversy but is widely considered to be due to its accumulation in lysosomes and subsequent disruption of their function (Kaufmann et al., 2009). Lysosomes have emerged as organelles with critical signaling roles regulating many aspects of cellular function (reviewed by Xu and Ren, 2015). They also play central roles in autophagy, the process by which cellular components are recycled (reviewed in Parzych and Klionsky, 2014). Chloroquine’s effects on lysosomes are likely to have effects on diverse downstream cellular processes including receptor signaling, with anti-inflammatory consequences (Wallace et al., 2012), and autophagy, with anti-tumor effects (Levy et al., 2017). There may therefore be multiple effects of chloroquine on hair cell function with consequences to their survival.

While both zebrafish and mouse cochlear hair cells are sensitive to chloroquine, there are differences in the nature of the dose-response functions. Loss of zebrafish hair cells approached an asymptote of about 40% reduction without further loss at higher concentrations of chloroquine, while basal outer hair cells in mouse cochlear cultures approach 100% loss with increasing dose. The underlying causes of asymptotic nature of the zebrafish dose-response function is unknown. We hypothesized that chloroquine may reduce its uptake and toxicity at higher concentrations by blocking hair cell mechanotransduction, a property required for the hair cell entry of some other compounds in zebrafish, including aminoglycosides and cisplatin (Thomas et al., 2012; Hailey et al. 2017). Although other quinoline loop derivatives have been shown to interfere with mechanotransduction (Farris et al., 2004; Ou et al., 2012), we found little effect of chloroquine on mechanotransduction-dependent FM1-43 dye uptake in zebrafish. Moreover we found that interfering with mechanotransduction had little effect on chloroquine toxicity. We suggest that there may be other reasons for incomplete toxicity in zebrafish, such as effective sequestration or efflux mechanisms that engage at higher chloroquine concentrations to protect hair cells.

Further research is needed to fully investigate mechanisms of hair cell death in the presence of chloroquine and more broadly how this class of drugs affect patients in their everyday lives. In addition to potential side effects directly detrimental to auditory perception and vestibular function, changes in these perceptual outcomes can create an increase in anxiety, depression, and fatigue resulting in a decrease in quality of life (Coelho and Balaban, 2015; Gomaa et al., 2014; Hornsby et al., 2016; Mira, 2008; Ohlenforst et al., 2017). Given the interest of chloroquine as a therapy for viral infections including Zika and SARS coronaviruses (Al-Bari, 2017), it will be important to consider monitoring auditory and vestibular function as these drugs are adopted to new therapies.

## Acknowledgments

This work was supported by the National Institutes of Deafness and other Communication Disorders R01DC005987, and Auditory Neuroscience Training Grant NIH T32DC005361.

## Author contributions

Work was conceptualized by SND, JAS, EWR and DWR; Investigation carried out by SND, PW and EDC; Manuscript written and edited by SND, EWR and DWR.

## References

Al-Bari, A.A., 2015. Chloroquine analogues in drug discovery: New directions of uses, mechanisms of actions and toxic manifestations from malaria to multifarious diseases. J. Antimicrob. Chemother. 70, 1608–1621. https://doi.org/10.1093/jac/dkv018

Bak-Coleman, J., Court, A., Paley, D.A., Coombs, S., 2013. The spatiotemporal dynamics of rheotactic behavior depends on flow speed and available sensory information. J. Exp. Biol. 216, 4011–4024. https://doi.org/10.1242/jeb.090480

Bernard, P., 1985. Alterations of Auditory Evoked Potentials during the Course of Chloroquine Treatment. Acta Otolaryngol. 99, 387–392.

Bogaczewicz, A., Sobów, T., 2017. Psychiatric adverse effects of chloroquine. Psychiatr. i Psychol. Klin. 17, 111–114. https://doi.org/10.15557/PiPK.2017.0012

Chiu, L.L., Cunningham, L.L., Raible, D.W., Rubel, E.W., Ou, H.C., 2008. Using the zebrafish lateral line to screen for ototoxicity. JARO - J. Assoc. Res. Otolaryngol. 9, 178–190. https://doi.org/10.1007/s10162-008-0118-y

Chowdhury, S., Owens, K.N., Herr, R.J., Jiang, Q., Chen, X., Johnson, G., Groppi, V.E., Raible, D.W., Rubel, E.W., Simon, J.A., 2018. Phenotypic Optimization of Urea-Thiophene Carboxamides to Yield Potent, Well Tolerated, and Orally Active Protective Agents against Aminoglycoside-Induced Hearing Loss. J. Med. Chem. 61, 84–97. https://doi.org/10.1021/acs.jmedchem.7b00932

Coelho, C.M., Balaban, C.D., 2015. Visuo-vestibular contributions to anxiety and fear. Neurosci. Biobehav. Rev. https://doi.org/10.1016/j.neubiorev.2014.10.023

Cortegiani, A., Ingoglia, G., Ippolito, M., Giarratano, A., Einav, S., 2020. A systematic review on the efficacy and safety of chloroquine for the treatment of COVID-19. J. Crit. Care 3–7. https://doi.org/10.1016/j.jcrc.2020.03.005

Farris, H.E., LeBlanc, C.L., Goswami, J., Ricci, A.J., 2004. Probing the pore of the auditory hair cell mechanotransducer channel in turtle. J. Physiol. 558, 769–792. https://doi.org/10.1113/jphysiol.2004.061267

Gomaa, M.A.M., Elmagd, M.H.A., Elbadry, M.M., Kader, R.M.A., 2014. Depression, Anxiety and Stress Scale in patients with tinnitus and hearing loss. Eur. Arch. Oto-Rhino-Laryngology. https://doi.org/10.1007/s00405-013-2715-6

Hailey, D.W., Esterberg, R., Limbo, T.H., Rubel, E.W., Raible, D.W., 2017. Fluorescent aminoglycosides reveal intracellular trafficking routes in mechanosensory hair cells The Journal of Clinical Investigation. J Clin Invest 127, 472–486. https://doi.org/10.1172/JCI85052

Harris, J.A., Cheng, A.G., Cunningham, L.L., MacDonald, G., Raible, D.W., Rubel, E.W., 2003. Neomycin-induced hair cell death and rapid regeneration in the lateral line of zebrafish (Danio rerio). JARO - J. Assoc. Res. Otolaryngol. 4, 219–234. https://doi.org/10.1007/s10162-002-3022-x

Hart, C.W., Naunton, R.F., 1964. The Ototoxicity of Chloroquine Phosphate. Arch. Otolaryngol. 80, 407–412.

Hennebert, D., Fernández, C., 1959. Ototoxicity of Quinine in Experimental Animals. AMA. Arch. Otolaryngol. 70, 321–333. https://doi.org/10.1001/archotol.1959.00730040329006

Hirose, Y., Simon, J.A., Ou, H.C., 2011. Hair cell toxicity in anti-cancer drugs: Evaluating an anticancer drug library for independent and synergistic toxic effects on hair cells using the zebrafish lateral line. JARO - J. Assoc. Res. Otolaryngol. 12, 719–728. https://doi.org/10.1007/s10162-011-0278-z

Hoppe, H.C., Van Schalkwyk, D.A., Wiehart, U.I.M., Meredith, S.A., Egan, J., Weber, B.W., 2004. Antimalarial quinolines and artemisinin inhibit endocytosis in Plasmodium falciparum. Antimicrob. Agents Chemother. https://doi.org/10.1128/AAC.48.7.2370-2378.2004

Hornsby, B.W.Y., Naylor, G., Bess, F.H., 2016. A taxonomy of fatigue concepts and their relation to hearing loss, in: Ear and Hearing. https://doi.org/10.1097/AUD.0000000000000289

Kapishnikov, S., Staalsø, T., Yang, Y., Lee, J., Pérez-Berná, A.J., Pereiro, E., Yang, Y., Werner, S., Guttmann, P., Leiserowitz, L., Als-Nielsen, J., 2019. Mode of action of quinoline antimalarial drugs in red blood cells infected by Plasmodium falciparum revealed in vivo. Proc. Natl. Acad. Sci. U. S. A. https://doi.org/10.1073/pnas.1910123116

Karlsson, K.K., Hellgren, U., Alvàn, G., Rombo, L., 1990. Audiometry as a possible indicator of quinine plasma concentration during treatment of malaria. Trans. R. Soc. Trop. Med. Hyg. 84, 765–767. https://doi.org/10.1016/0035-9203(90)90069-Q

Kaufmann, A.M., Goldman, S.D.B., Krise, J.P., 2009. A fluorescence resonance energy transfer-based approach for investigating late endosome-lysosome retrograde fusion events. Anal. Biochem. 386, 91–97. https://doi.org/10.1016/j.ab.2008.11.036

Kirkwood, N.K., O’Reilly, M., Derudas, M., Kenyon, E.J., Huckvale, R., Van Netten, S.M., Ward, S.E., Richardson, G.P., Kros, C.J., 2017. d-tubocurarine and berbamine: Alkaloids that are permeant blockers of the hair cell’s mechano-electrical transducer channel and protect from aminoglycoside toxicity. Front. Cell. Neurosci. 11, 1–15. https://doi.org/10.3389/fncel.2017.00262

Kitcher, S.R., Kirkwood, N.K., Camci, E.D., Wu, P., Gibson, R.M., Redila, V.A., Simon, J.A., Rubel, E.W., Raible, D.W., Richardson, G.P., Kros, C.J., 2019. ORC-13661 protects sensory hair cells from aminoglycoside and cisplatin ototoxicity. JCI Insight 4. https://doi.org/10.1172/jci.insight.126764

Kotecha, B., Richardson, G.P., 1994. Ototoxicity in vitro: effects of neomycin, gentamicin, dihydrostreptomycin, amikacin, spectinomycin, neamine, spermine and poly-l-lysine. Hear. Res. https://doi.org/10.1016/0378-5955(94)90232-1

Levy, J.M.M., Towers, C.G., Thorburn, A., 2017. Targeting autophagy in cancer. Nat. Rev. Cancer. https://doi.org/10.1038/nrc.2017.53

Matz, G.J., Naunton, R.F., 1968. Ototoxicity of Chloroquine. Arch. Otolaryngol. 88, 370–372. https://doi.org/10.1017/s0022215100085960

Mira, E., 2008. Improving the quality of life in patients with vestibular disorders: The role of medical treatments and physical rehabilitation. Int. J. Clin. Pract. https://doi.org/10.1111/j.1742-1241.2006.01091.x

Mukherjee, D.K., 1979. Chloroquine ototoxicity—a reversible phenomenon? J. Laryngol. Otol. 93, 809–815.

Mukherjee, D.K., Mukherjee, K., 1979. Ototoxicity of Commonly Used Pharmaceutical Preparations. Niger. Med. J. 9, 52–57.

Nielsen-Abbring, F.W., Perenboom, R.M., van der Hulst, R.J.A.M., 1990. Quinine-Induced Hearing Loss. J. Oto-rhino-laryngology its Borderl. 52, 65–68.

Ohlenforst, B., Zekveld, A.A., Jansma, E.P., Wang, Y., Naylor, G., Lorens, A., Lunner, T., Kramer, S.E., 2017. Effects of hearing impairment and hearing aid amplification on listening effort: A systematic review. Ear Hear. 38, 267–281. https://doi.org/10.1097/AUD.0000000000000396

Ou, H.C., Keating, S., Wu, P., Simon, J.A., Raible, D.W., Rubel, E.W., 2012. Quinoline ring derivatives protect against aminoglycoside-induced hair cell death in the zebrafish lateral line. JARO - J. Assoc. Res. Otolaryngol. 13, 759–770. https://doi.org/10.1007/s10162-012-0353-0

Owens, K.N., Santos, F., Roberts, B., Linbo, T., Coffin, A.B., Knisely, A.J., Simon, J.A., Rubel, E.W., Raible, D.W., 2008. Identification of genetic and chemical modulators of zebrafish mechanosensory hair cell death. PLoS Genet. 4. https://doi.org/10.1371/journal.pgen.1000020

Parzych, K.R., Klionsky, D.J., 2014. An overview of autophagy: Morphology, mechanism, and regulation. Antioxidants Redox Signal. https://doi.org/10.1089/ars.2013.5371

Phillips-Howard, P.A., ter Kuile, F.O., 1995. CNS Adverse Events Associated with Antimalarial Agents. Drug Saf. 12, 370–383.

Pickett, S.B., Thomas, E.D., Sebe, J.Y., Linbo, T., Esterberg, R., Hailey, D.W., Raible, D.W., 2018. Cumulative mitochondrial activity correlates with ototoxin susceptibility in zebrafish mechanosensory hair cells. Elife 7, 1–25. https://doi.org/10.7554/eLife.38062

Plantone, D., Koudriavtseva, T., 2018. Current and Future Use of Chloroquine and Hydroxychloroquine in Infectious, Immune, Neoplastic, and Neurological Diseases: A Mini-Review. Clin. Drug Investig. https://doi.org/10.1007/s40261-018-0656-y

Richardson, G.P., Russell, I.J., 1991. Cochlear cultures as a model system for studying aminoglycoside induced ototoxicity. Hear. Res. https://doi.org/10.1016/0378-5955(91)90062-E

Roche, R.J., Silamut, K., Pukrittayakamee, S., Looareesuwan, S., Molunto, P., Boonamrung, S., White, N.J., 1990. Quinine induces reversible high-tone hearing loss. Br. J. Clin. Pharmacol. 29, 780–782.

Rosner, B., 2011. Hypothesis testing: two-sample inference–estimation of sample size and power for comparing two means. Fundam. Biostat. 7th ed., Brooks/Cole, Cengage Learn.

Rüsch, A., Kros, C.J., Richardson, G.P., 1994. Block by amiloride and its derivatives of mechano-electrical transduction in outer hair cells of mouse cochlear cultures. J. Physiol. 474, 75–86. https://doi.org/10.1113/jphysiol.1994.sp020004

Russell, I.J., Richardson, G.P., 1987. The morphology and physiology of hair cells in organotypic cultures of the mouse cochlea. Hear. Res. 31, 9–24. https://doi.org/10.1016/0378-5955(87)90210-3

Sahraei, Z., Shabani, M., Shokouhi, S., Saffaei, A., 2020. Aminoquinolines Against Coronavirus Disease 2019 (COVID-19): Chloroquine or Hydroxychloroquine. Int. J. Antimicrob. Agents 2019, 105945. https://doi.org/10.1016/j.ijantimicag.2020.105945

Santos, F., MacDonald, G., Rubel, E.W., Raible, D.W., 2006. Lateral line hair cell maturation is a determinant of aminoglycoside susceptibility in zebrafish (Danio rerio). Hear. Res. 213, 25–33. https://doi.org/10.1016/j.heares.2005.12.009

Scherbel, A.L., Harrision, J.W., Atdjian, M., 1958. Further observations on the use of 4-aminoquinoline compounds in patients with rheumatoid arthritis or related diseases. Cleve. Clin. Q. 25, 95–111.

Shrivastav, A., Singh, S., Agarwal, M., 2016. Chloroquine Retinopathy. Delhi Ophthalmol. Soc. 22, 19–22. https://doi.org/10.1016/0002-9394(62)93396-2

Thomas, E.D., Cruz, I.A., Hailey, D.W., Raible, D.W., 2015. There and back again: Development and regeneration of the zebrafish lateral line system. Wiley Interdiscip. Rev. Dev. Biol. 4, 1–16. https://doi.org/10.1002/wdev.160

Toone, E.C., Hayden, D., Ellman, H.M., 1965. Ototoxicity of Chloroquine. Proc. Annu. Meet. Am. Rheum. Assoc. 475–476.

U.S. Food and Drug Administration, 2019. FDA Adverse Events Reporting System (FAERS) Public Dashboard [WWW Document]. URL https://fis.fda.gov/sense/app/d10be6bb-494e-4cd2-82e4-0135608ddc13/sheet/8eef7d83-7945-4091-b349-e5c41ed49f99/state/analysis (accessed 1.6.20).

Wallace, D.J., Gudsoorkar, V.S., Weisman, M.H., Venuturupalli, S.R., 2012. New insights into mechanisms of therapeutic effects of antimalarial agents in SLE. Nat. Rev. Rheumatol. https://doi.org/10.1038/nrrheum.2012.106

Xu, H., Ren, D., 2015. Lysosomal Physiology. Annu. Rev. Physiol. https://doi.org/10.1146/annurev-physiol-021014-071649

Zhou, D., Dai, S.-M., Tong, Q., 2020. COVID-19: a recommendation to examine the effect of hydroxychloroquine in preventing infection and progression. J. Antimicrob. Chemother. 4–7. https://doi.org/10.1093/jac/dkaa114

